# *Drosophila* antimicrobial peptides and lysozymes regulate gut microbiota composition and abundance

**DOI:** 10.1101/2021.03.19.436153

**Authors:** A. Marra, M.A. Hanson, S. Kondo, B. Erkosar, B. Lemaitre

## Abstract

The gut microbiota affects the physiology and metabolism of animals and its alteration can lead to diseases such as gut dysplasia or metabolic disorders. Several reports have shown that the immune system plays an important role in shaping both bacterial community composition and abundance in *Drosophila*, and that immune deficit, especially during aging, negatively affects microbiota richness and diversity. However, there has been little study at the effector level to demonstrate how immune pathways regulate the microbiota. A key set of *Drosophila* immune effectors are the antimicrobial peptides (AMPs), which confer defense upon systemic infection. AMPs and lysozymes, a group of digestive enzymes with antimicrobial properties, are expressed in the gut and are good candidates for microbiota regulation. Here, we take advantage of the model organism *Drosophila melanogaster* to investigate the role of AMPs and lysozymes in regulation of gut microbiota structure and diversity.

Using flies lacking AMPs and newly generated lysozyme mutants, we colonized gnotobiotic flies with a defined set of commensal bacteria and analyzed changes in microbiota composition and abundance in vertical transmission and aging contexts through 16S rRNA gene amplicon sequencing. Our study shows that AMPs and, to a lesser extent, lysozymes are necessary to regulate the total and relative abundance of bacteria in the gut microbiota. We also decouple the direct function of AMPs from the IMD signaling pathway that regulates AMPs but also many other processes, more narrowly defining the role of these effectors in the microbial dysbiosis observed in IMD-deficient flies upon aging.

**Importance:** This study advances current knowledge in the field of host-microbe interactions by demonstrating that the two families of immune effectors, antimicrobial peptides and lysozymes, actively regulate the gut microbiota composition and abundance. Consequences of the loss of these antimicrobial peptides and lysozymes are exacerbated during aging, and their loss contributes to increased microbiota abundance and shifted composition in old flies. This work shows that immune effectors, typically associated with resistance to pathogenic infections, also help shape the beneficial gut community, consistent with the idea that host-symbiont interactions use the same ‘language’ typically associated with pathogenesis.

## Introduction

The microbiota is the complex array of microbes commonly associated with the digestive tract of animals. This bacterial consortium greatly affects host physiology, for example by promoting immune function or intestinal homeostasis (1–4). Imbalance of the microbiota, called dysbiosis, has been identified as a cause of gut dysplasia and chronic inflammatory diseases, especially during aging (5).

The fruit fly *Drosophila melanogaster* is a powerful model to decipher host-microbe interactions (6–8). Its genetic tractability, the possibility to generate gnotobiotic animals and the simplicity of its natural microbiota have made *Drosophila melanogaster* a convenient model to gain insight into host-microbiota relationships (7, 9, 10). *Drosophila* harbors a simple gut microbiota composed of only a few dominant species, mainly belonging to *Acetobacteraceae* and *Lactobacillaceae*, which influence multiple aspects of fly physiology, such as growth (11, 12), behavior (13), lifespan (14) and infection resistance (15, 16). In turn, the microbiota can be shaped by various host and environmental factors such as food composition, or age of the flies (17–21).

Innate immunity is a key regulator of microbial abundance in *Drosophila* (22–25). Upon acute bacterial infection, *Drosophila* immune responsive tissues (the fat body and hemocytes in systemic infection, and epithelium in local infection) sense microbe-associated molecular patterns (MAMPs) to activate signaling pathways. In *Drosophila*, two immune pathways, the Immune deficiency (IMD) and Toll pathways, regulate the expression of genes encoding immune effectors that fight invading microbes (22, 26, 27). Studies in *Drosophila* have revealed a key role of the IMD pathway in the gut to fight pathogens and keep symbiotic bacteria in check (28–30). It is however unclear how the IMD pathway can effectively combat pathogens but tolerate symbiotic microbiota members in the digestive tract. In fact, the microbiota induces a low level of activation of the IMD pathway (28). Several reports have demonstrated that immune tolerance towards the indigenous microbiota is sustained by several negative feedback loops that prevent hyperactivation of the IMD pathway by peptidoglycan (the bacterial elicitor recognized by IMD pathway) released from commensal bacteria (31–33). Compartmentalization of the immune response to restricted areas can also favor microbiota growth and control (34, 35). However, IMD pathway activation is necessary to regulate both microbiota composition and proliferation, and dysregulation of this pathway leads to abnormal bacterial growth and premature death of the host (28, 32, 33, 36). Notably, mutations affecting the IMD transcription factor Relish lead to a higher gut microbiota load and a shifted bacterial composition compared to wild-type flies (28, 37). Moreover, aged *Relish* mutant flies display dysbiosis associated with a loss of gut epithelium integrity and premature death of the animals (21, 38). Collectively, these studies point to an important role of the IMD pathway in control of the microbiota, notably during aging. However, the IMD pathway regulates hundreds of immune effectors, and affects numerous physiological processes such as enterocyte delamination and digestion (39–43). As previous studies have used mutations that suppress the whole pathway (e.g *Relish*), the precise role of individual immune effectors downstream of the IMD pathway in shaping the gut microbial community has remained elusive.

Antimicrobial peptides (AMPs) are molecules that contribute to innate defenses by targeting the negatively charged membranes of microbes (27). These peptides are produced in large quantities by the fat body during systemic infection, but also in local epithelia such as the gut. Seven classic families of inducible AMPs with several isoforms have been identified in *D. melanogaster* (27, 44). Use of CRISPR/CAS9 has recently enabled the generation of individual and combined AMP mutants, allowing direct investigation of their role in host defense (45). Hanson et al. showed that *Drosophila* AMPs are essential for resisting infection by Gram-negative bacteria that trigger the IMD pathway, but appear to be less involved in defense against Gram-positive bacterial infection (45).

Another key group of effector proteins that are potential regulators of Gram-positive bacteria in the gut are the lysozymes (46, 47). Lysozymes specifically cleave peptidoglycan exposed on the cell wall of Gram-positive bacteria (48). The *Drosophila* genome encodes at least 17 putative lysozymes, whose functions have never been formally addressed. Among them, six lysozyme genes (*LysB, D, E, P, S*, and *X*) are clustered in the genome at cytogenetic map position 61F. This group of lysozyme genes, notably *LysB, LysD, LysE*, and *LysP*, is strongly expressed in the digestive tract (46), and may contribute to digestive activities of the gut by degrading peptidoglycan from dietary bacteria. Furthermore, lysozyme genes are expressed in the gut upon microbiota colonization in *Drosophila*, and these proteins have been proposed to modulate immune signaling (28, 31, 49). Lysozymes may contribute to gut immunity either as direct antimicrobials, or by cleaving peptidoglycan and modulating activation of the IMD pathway (31). As such, AMPs and lysozymes may shape microbiota composition by direct interactions with microbes.

In this study, we decipher the role of two classes of antimicrobial effectors of the *Drosophila* digestive tract, the antimicrobial peptides (AMPs) and the lysozymes, on the gut microbiota. We characterized the microbiota composition in mutant flies lacking either the 14 AMP genes from seven gene families or the four gut-specific lysozyme-encoding genes in a gnotobiotic setup using 16S rRNA gene amplicon sequencing (referred to as 16S sequencing). We also assessed the role of these effectors in controlling the abundance of individual microbiota members by performing mono-association experiments. Finally, we confirmed that certain immune effectors can directly control the proliferation of microbiota members by performing systemic infections. Our findings demonstrate a direct role for both AMPs and lysozymes in controlling both the composition and abundance of the microbiota in *Drosophila melanogaster*.

## Results

### Impact of AMPs and lysozymes on microbiota composition

To decipher the role of AMPs and lysozymes in the regulation of gut microbiota composition, we performed 16S sequencing on gnotobiotic flies. DrosDel isogenic flies with the following genotypes were used for all experiments: the wild-type strain *w*^*1118*^ (referred to as *w*), a compound mutant strain lacking *Defensin, Cecropins (*4 *genes), Drosocin, Diptericins (*2 *genes), Attacins (*4 *genes), Metchnikowin* and *Drosomycin*, referred to as *ΔAMPs*^*14*^ (Carboni et al., in preparation), and a newly-generated lysozyme-deficient mutant (referred to *LysB-P*^Δ^). The *LysB-P*^Δ^ mutation is an 11.5kb deletion, removing *LysC* (a putative pseudogene) and the four lysozyme genes (i.e. *Lys B, LysD, LysE, Lys P*) that are known to be strongly expressed in the digestive tract (46) (**Fig S1**). As expected, gut extracts from *LysB-P*^Δ^ flies have reduced lysozyme activity *ex vivo* as monitored by their ability to digest peptidoglycan from *E. faecalis* (**Fig. S1**). We additionally included *Relish* (*Rel*^*E20*^) flies lacking IMD signaling as a comparative control to determine to what extent AMPs contribute to the phenotype of IMD-deficient flies. To avoid pre-existing microbial community biases in different fly stocks, we performed this analysis in a gnotobiotic system with two different experimental designs. First, we analyzed the microbiota of 12-day old flies with gut bacteria acquired through vertical transmission from gnotobiotic parental flies (i.e. germ-free parents inoculated with a known community upon adult emergence) (**Fig 1A**). Second, we analyzed aging-dependent changes in the adult microbiota. Here, we inoculated emerging germ-free (referred as GF) adults with a known microbiota and analyzed changes in the community structure 10 and 29 days after colonization (**Fig 2A**). In this way, we uncoupled the effects of juvenile development and metamorphosis from the adult microbiota composition and abundance.

**Fig 1.**
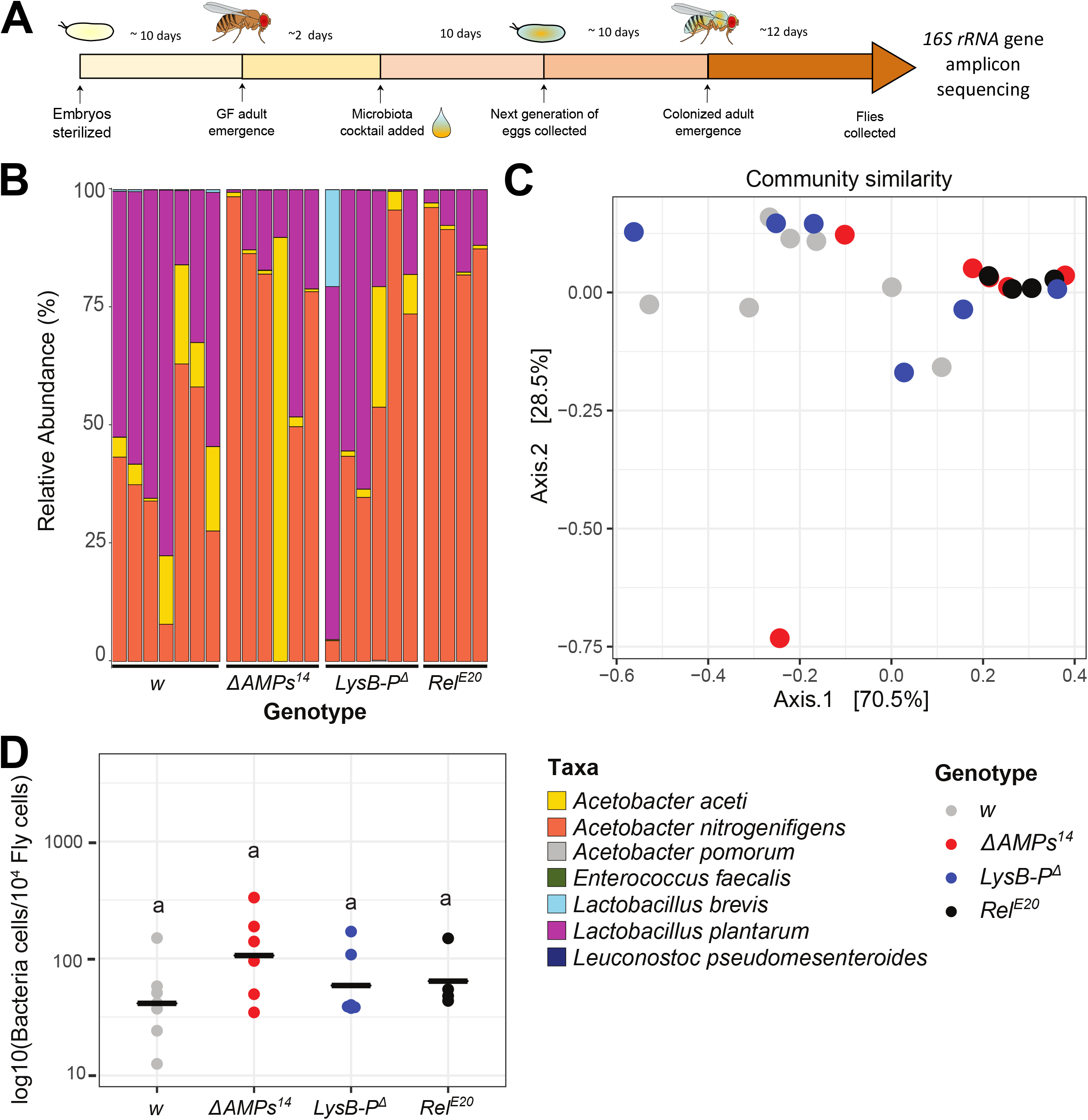
The role of AMPs and lysozymes on microbiota composition and abundance in a gnotobiotic vertical transmission setup. A) Scheme of the experimental procedure for fly colonization and collection for 16S rRNA gene amplicon sequencing. Parental embryos were collected, sterilized in 3% bleach and kept on antibiotic food until the adult stage. Emerging GF flies were then associated with a bacterial cocktail (microbiota cocktail) containing six representative microbiota members. Their eggs were collected over 3 days, allowed to develop to adulthood, and finally the microbiota of their adult female progeny was analyzed ~12 days after emergence. B) Relative community composition of the gut microbiota in wild-type iso *w*^*1118*^ (*w*) wild-type flies, *Relish* (*Rel*^*E20*^), antimicrobial peptide (*ΔAMPs*^*14*^), and gut lysozyme (*LysB-P*^*Δ*^) mutants as determined by 16S rRNA gene amplicon sequencing. Each bar represents a biological replicate of multiple pooled flies (see **Table S1** numbers of flies included in each sample). C) Principal coordinate analysis (PCoA) of gut communities in *w* wild-type flies, *Rel*^*E20*^, *ΔAMPs*^*14*^ and *LysB-P*^*Δ*^ as determined by 16S rRNA gene amplicon sequencing. Overall colocalization of *ΔAMPs*^*14*^ (red dots) and *Rel*^*E20*^ (black dots) samples and separation of these from wild-type (grey dots) samples shows that *ΔAMPs*^*14*^ and *Rel*^*E20*^ samples are similar to each other and differ from wild-type samples. Stochastic distribution of *LysB-P*^*Δ*^ samples shows high variability in community structures between samples. D) Absolute quantification by qPCR of the total number of bacterial cells normalized to the host gene *Actin5C*. Horizontal black bars show mean values. Details of the statistical outcomes are provided in **Supplementary Table S2**.

**Fig 2.**
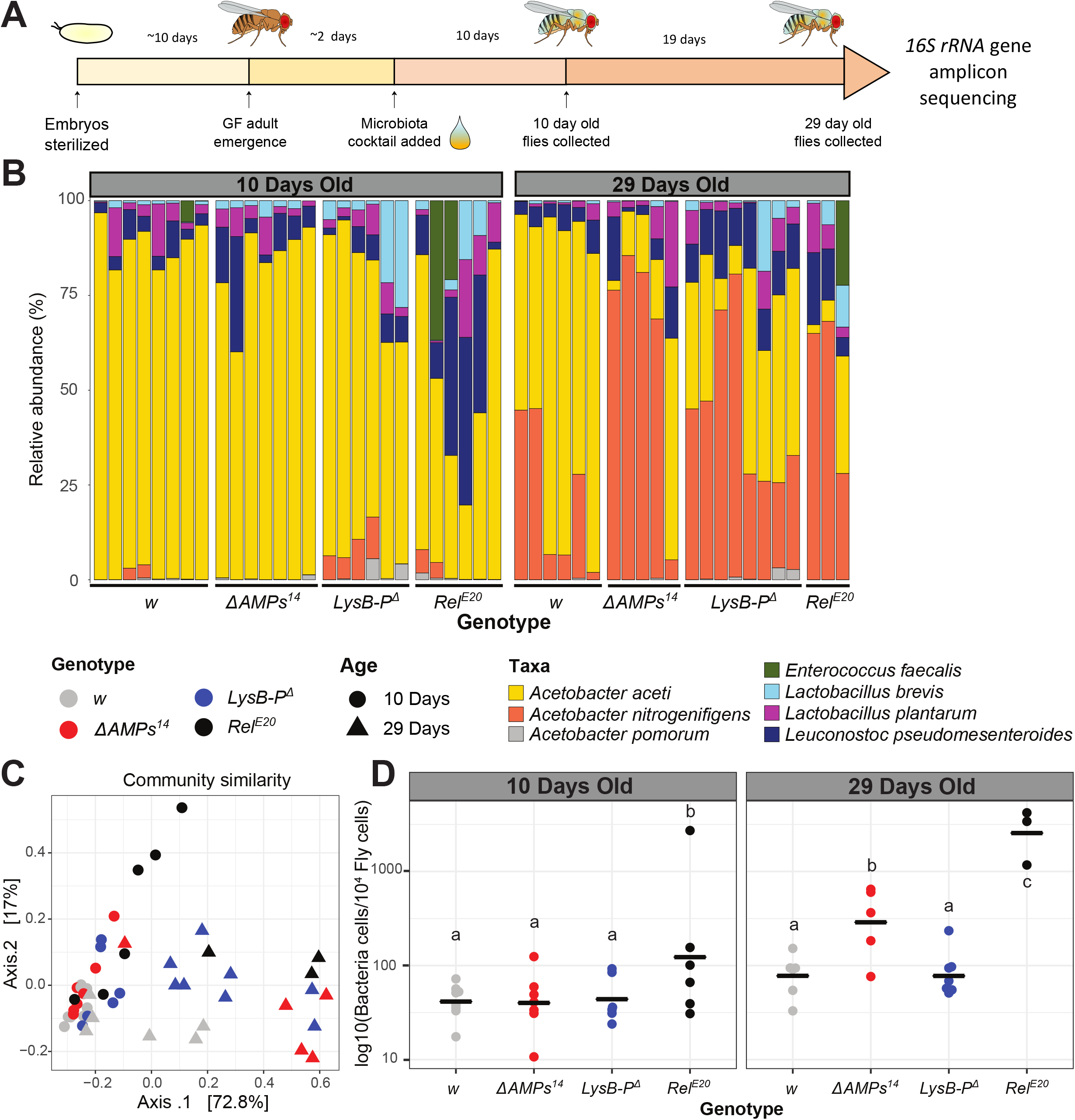
The role of AMPs and lysozymes in microbiota composition and abundance on adult microbiota in a gnotobiotic setup. A) Scheme of the experimental procedure for fly colonization and collection for 16S rRNA gene amplicon sequencing. Embryos were collected, sterilized in 3% bleach, and kept on antibiotic food until the adult stage. Emerging GF flies were associated with a bacterial cocktail containing six representative microbiota members. Females were collected for DNA extraction and 16S rRNA gene amplicon sequencing 10 and 29 days after colonization. See **Table S1** for the number of flies included in each sample. B) Relative community composition of the gut microbiota in iso *w*^*1118*^ (*w*) wild-type flies and *Relish* (*Rel*^*E20*^), antimicrobial peptide (*ΔAMPs*^*14*^), and gut lysozyme (*LysB-P*^*Δ*^) mutants 10 days (left panel) and 29 days (right panel) after colonization. Each bar in the plot represents a biological replicate with a pool of 5 flies each. C) Principal coordinate analysis based on Bray–Curtis dissimilarities on the gut communities of *w* control flies, *Rel*^*E20*^, *ΔAMPs*^*14*^, and *LysB-P*^*Δ*^ mutants 10 and 29 days after colonization based on 16S rRNA gene amplicon sequencing. Separation of the 10-day old (dots) and 29-day old clusters on the first axis, indicates that aging is the major factor defining bacterial community composition in adults. Separation *ΔAMPs*^*14*^ and *Rel*^*E20*^ (red and black triangles) from wild-type and *LysB-P*^*Δ*^ (grey and blue triangles) on the same axis in 29-day samples indicates that aging and loss of immune effectors act on microbiota composition in similar directions. D) Absolute quantification of the total number of bacterial cells by qPCR, normalized to the host gene *Actin5C*. Horizontal black bars show mean values. Details of the statistical outcomes are provided in Supplementary **Table S2**.

We inoculated the flies with a cocktail of six bacterial isolates that were previously described as common *Drosophila* microbiota members (19), or that were associated with the food that was used in this study (see Methods) (8, 10, 19). These included previously characterized bacterial species as members of the *Drosophila* gut microbiota: *Acetobacter pomorum* (50), *Lactobacillus plantarum* (11), and *Enterococcus faecalis* (51). Our cocktail also included some incompletely characterized bacterial strains: an *Acetobacter sp*. (52), an isolate of *Lactobacillus brevis* and an isolate of *Leuconostoc pseudomesenteroides* (see Materials and Methods, Supplementary text 1). *Acetobacter*, a genus of Gram-negative bacteria, and *Lactobacillus plantarum*, a Gram-positive species, both have DAP-type peptidoglycan known to activate the IMD pathway (51, 53–55). In contrast, *Leuconostoc pseudomesenteroides, Lactobacillus brevis* (53, 56) and *Enterococcus faecalis* are Gram-positive bacteria with Lysine-type peptidoglycan (57), which typically activates the Toll pathway during systemic infections. Although there is no evidence for a role of the Toll pathway in the midgut (22, 43, 54), Lys-type Gram-positive bacteria can induce a basal immune reaction in the gut through the release of the metabolite uracil, which activates ROS production through the Duox enzyme (58). 16Ssequencing of the six-component cocktail yielded 7 Amplicon Sequence Variants (ASVs (59) are also referred to as zero-noise OTUs (60) or sub-OTUS (61)) across the 72 samples, with a minimum of 42’596 reads per sample after quality and abundance filtering (see Methods and **Supplementary Table S1** for details). These ASVs mapped to the known species in the inoculum cocktail. Sequencing showed that the *Acetobacter* sp. (52) fraction mapped to two ASVs that were distinguishable by a single nucleotide difference in their 16S amplicon. These ASVs were associated with two closely related species, *A. aceti* and *A. nitrogenifigens* (62, 63), based on their highly similar sequence.

We first focused our analysis on flies with microbiota acquired through vertical transmission from parents raised in a gnotobiotic environment (**Fig 1A**, see Methods). We found that *Rel*^*E20*^ and *ΔAMPs*^*14*^ flies harbored communities dominated by *A. nitrogenifigens*, whereas the wild-type strain had a greater prevalence of *La. plantarum* (**Fig 1B**). In contrast, *LysB-P*^*Δ*^ flies had highly variable community compositions (see below), suggesting a different mode of action for these genes compared to the AMPs (**Fig 1B**).

Similarities between bacterial communities were assessed using β-diversity analyses. Dissimilarities between all samples were calculated using Bray-Curtis distances plotted in a multi-dimensional space using Principal Component Analysis (PCoA). This was complemented with an analysis of the dispersal (variability and spread) of the communities, and a permutation based, multivariate analysis of variance was applied to test statistical significance. These analyses showed that community compositions within *LysB-P*^*Δ*^ and *ΔAMPs*^*14*^ sample groups were more variable than the wild-type (**Fig S2A**, 0.05<p<0.1), in that communities of some samples resembled wild*-*type flies, while others resembled *Rel*^*E20*^ flies (**Fig 1C**). One *LysB-P*^*Δ*^ sample had a completely different profile to all other samples, with higher abundance of *La. brevis* (**Fig 1B**). This suggests that the loss of AMPs or lysozymes increases stochasticity in microbiota composition. Surprisingly, communities in *Rel*^*E20*^ mutants were more consistent between replicates, which indicates either that the stochasticity is not due to perturbation of the immune response or that the communities in these mutants stabilize earlier than in other genotypes due to other factors regulated by the IMD pathway.

In terms of community composition, distribution of data in the PCoA shows that *ΔAMPs*^*14*^ samples mimic *Rel*^*E20*^ (Pairwise ADONIS: p-adjustedΔAMPs vs Rel=0.5), and both differ noticeably from the wild-type, as demonstrated by general colocalization of *ΔAMPs*^*14*^ and *Rel*^*E20*^ samples, and separation from the wild-type samples (**Fig 1C**, Pairwise ADONIS: p-adjustedw vs Rel=0.02, p-adjustedw vs ΔAMPs=0.06). This suggests that loss of AMPs recapitulates the effect of a general loss of the IMD pathway on the microbiota structure. As expected from the variable community composition found in *LysB-P*^*Δ*^ mutants (Fig 1B), the PCoA did not reveal a distinct cluster for these samples (**Fig 1C**).

Finally, we measured total bacterial loads in our samples using universal *16S rRNA* gene primers (64) and *Drosophila Actin 5C* primers (52). We did not detect a statistically significant difference in total 16S rRNA gene copy numbers between the different genotypes, indicating that wild-type and mutant flies do not harbor different quantities of microbes in these conditions (**Fig 1D**).

Overall, our results show that the microbiota composition in *ΔAMPs*^*14*^ flies is similar to the microbiota of *Rel*^*E20*^ mutants that completely lack IMD signaling, suggesting that the changes in community composition observed in IMD pathway mutants is at least partly due to the specific loss of AMP production.

### Control of microbiota structure by AMPs and lysozymes during aging

Next, we focused on the microbiota structure of adult flies that were raised in GF conditions throughout larval development and colonized only after emergence. We analyzed microbiota of these flies at both 10 and 29 days after colonization (**Fig 2A**). Here, microbial communities were generally dominated by the two *Acetobacter* variants. 10 days after colonization, *A. aceti* was the most abundant species, whereas by 29 days after colonization, *A. nitrogenifigens* was the dominant species., suggesting distinct competitive ability of the two bacteria tied to the 16S sequence variants detected in our *Acetobacter sp*. isolate.

As *Rel*^*E20*^ mutants died earlier than other genotypes during the aging process, only three samples with fewer flies than other genotypes were included for the 29-day time point (**Fig 2B, Table S1**). *Rel*^*E20*^ mutants harbored elevated abundance of *E. faecalis* in 1/3 of the samples, which was not observed in other genotypes. Some samples in this genotype also had higher proportions of *La. plantarum* and *Le. pseudomesenteroides* (in two and three samples respectively) at day 10, a trend that was not observed at day 29 (**Fig 2B**). However, we cannot conclude whether this change in community structure is real or a consequence of high mortality in this genotype leading to analysis of the survivors only. In contrast to the vertical transmission setup (**Fig S2A**), *Rel*^*E20*^ communities had high dispersal: highest variation was observed at 10 days (**Fig S2B**), and decreased at 29 days (**Fig S2C**). This indicates that immunity mutations cause stochasticity in microbiota composition, but the communities are still capable of stabilizing over a long period of time.

β-diversity and PCoA clearly shows significant (ADONIS p=0.001) separation of the 10-day old and 29-day old flies on the first axis, clearly pointing to aging as the major factor defining the community composition in adults (**Fig 2C**). Interestingly, in 29-day old flies, *ΔAMPs*^*14*^ and *Rel*^*E20*^ were separated from wild-type and *LysB-P*^*Δ*^ mutants on the same axis (Fig 2C), indicating that aging and loss of immune effectors act on microbiota composition in similar directions. In 10-day old flies, we did not see similar clustering of samples except for *Rel*^*E20*^ mutants, which were more widely dispersed on the plot (**Fig 2C**). This indicates that mutations in IMD pathway act on microbiota composition differently in young versus old flies. A statistically significant Genotype x Age interaction (ADONIS p=0.03) supports this interpretation.

Careful examination of the relative abundance of bacteria in wild-type and mutant flies reveals interesting trends (**Fig 2B**). We found that wild-type flies maintained *Acetobacter aceti* as the dominant *Acetobacter* ASV even after 29 days, while the proportion of lactobacilli in the community remained small. However, although *Acetobacter aceti* was similarly abundant at 10 days *ΔAMPs*^*14*^ and *LysB-P*^*Δ*^ flies, *Acetobacter nitrogenifigens* became predominant in 29-day samples, and the proportion of lactobacilli in some samples was higher compared to wild-type, particularly in *LysB-P*^*Δ*^ flies. This change in relative abundances was even more dramatic in *Rel*^*E20*^ mutants, which were distinguished by disproportionate loads of *Acetobacter nitrogenifigens* and lactobacilli.

Investigation of each time point separately showed that loss of AMPs did not affect the community composition in 10-day old flies (**Fig 2B**, Pairwise ADONIS, qw vs ΔAMPs=0.1). However, loss of lysozymes had detectable effects (Pairwise ADONIS, qw vs Lys=0.02) on the abundance of *A. pomorum, A. nitrogenifigens*, or *La. brevis* depending on the samples, which further supports the idea of increased stochasticity in *LysB-P*^Δ^ mutants compared to wild-types. This stochasticity is clearly shown by the community dispersal (**Fig S2B**).

At 29 days, microbial communities in the wild-type differed from those of *ΔAMPs*^*14*^, *LysB-P*^Δ^ and *Rel*^*E20*^ genotypes (Pairwise ADONIS p-adjustedw vs ΔAMPs =0.04, p-adjustedw vs Lys= 0.04, p-adjustedw vs *Rel* = 0.04, **Fig 2B, 2C**). In *ΔAMPs*^*14*^ the relative abundance of Gram-negative *A*.

*nitrogenifigens* consistently increased, whereas in *LysB-P*^*Δ*^ mutants the relative abundance of Gram-positive lactobacilli increased (**Fig 2B**). This suggests that lysozymes act preferentially on Gram-positive bacteria, and the action of AMPs is limited to *Acetobacteraceae*. As all genotypes contain communities that are similarly variable (**Fig S2C**), the observed differences in community composition at day 29 are unlikely to be an artefact of heterogeneity in variance among different groups.

Analysis of the total microbiota abundance showed that bacterial load differed between genotypes mainly in aged flies. At 10 days old, *Rel*^*E20*^ flies harbored significantly higher amounts of total bacteria compared to the other genotypes, primarily due to one sample that had a high load typical in 29-day old samples of this genotype (**Fig 2D**). In 29-day old flies both *ΔAMPs*^*14*^ and particularly *Rel*^*E20*^ flies had higher bacterial loads (**Fig 2D**). These data support the notion that the IMD pathway is crucial in regulating microbiota load as the flies age and that AMPs significantly contribute to this effect of the IMD pathway.

In agreement with previous reports, our data show that microbial community composition shifts and bacterial load increases with age (17, 18, 28), and that this effect is exacerbated by loss of antimicrobials.

### Effect of AMPs and lysozymes on individual microbiota members

16S sequencing provided us a first glimpse of how AMPs and gut lysozymes regulate microbiota structure at the community level. To further characterize the effect of these antimicrobials on individual microbiota members, we used a mono-association setup where we colonized flies with each bacterial isolate from the commensal cocktail used in the 16S sequencing experiment. GF adult females were mono-associated with a single bacterial species and their load was measured 6 days after colonization by qPCR (**Fig 3A**). We quantified 16S rRNA gene copies using primers that recognize *Acetobacteraceae* (65) and Firmicutes (including *La. plantarum, Le. pseudomesenteroides* and *La. brevis)* and normalized their abundance to host cells using primers for *Actin 5C* (52) (**Fig 3B**).

**Fig 3.**
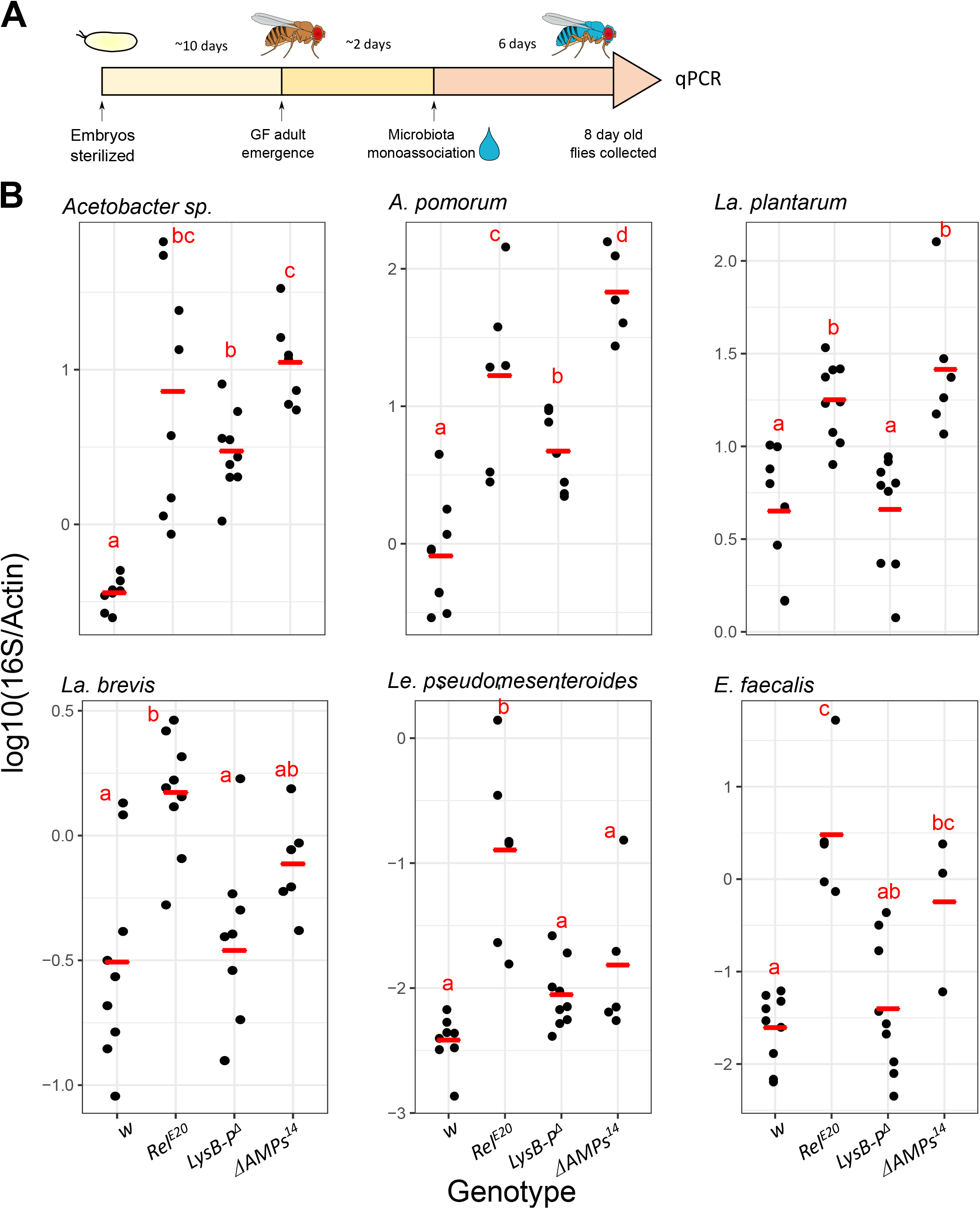
Regulation of individual microbiota members in mono-association. A) Scheme of the experimental procedure of the mono-association experiment. Embryos were collected, sterilized in 3% bleach, and kept on antibiotic food until the adult stage. Newly emerged GF flies were then mono-associated with a single bacterial isolate. Six days after colonization, the host and bacterial DNA was extracted and qPCR analysis of the microbial load was performed. B-G) Total microbial load was determined by quantitative PCR (qPCR) in female flies 6 days after mono-association. *iso w*^*1118*^ (*w*) wild-type flies and *Relish* (*Rel*^*E20*^), antimicrobial peptide (*ΔAMPs*^*14*^), and gut lysozyme (*LysB-P*^*Δ*^) mutant flies were included. Bacterial loads were assessed by qPCR with family/phylum specific 16S rRNA gene primers and normalized to the host gene *Actin5C*. Red horizontal bars show mean values. Each dot represents a sample containing five individuals. Letters represent statistical significance (p<0.05) of adjusted p-values (FDR) from pairwise contrasts obtained from a main general linear mixed model; samples with shared letters are not statistically different from each other.

As expected, all mono-associated taxa established a higher load in *Rel*^*E20*^ flies compared to wild-type flies (**Fig 3B**). Interestingly, abundance of both *Acetobacter sp*. and *A. pomorum* isolates were high in *LysB-P*^*Δ*^ but especially in *ΔAMPs*^*14*^ mutants (**Fig 3B**), indicating that AMPs most prominently control the proliferation of these Gram-negative microbiota members. Surprisingly, in contrast to shifts towards increased lactobacilli seen in the absence of lysozymes in gnotobiotic experiments (**Fig 1, Fig 2**), mono-associated *La. plantarum* increased in abundance in the absence of AMPs but not lysozymes (**Fig 3B**). This was surprising considering that lysozymes are expected to digest Gram-positive bacteria. The differing trends resulting from these approaches may depend on bacterial community dynamics in gnotobiotic experiments, or age-related differences between the experimental setups. Interestingly, *ΔAMPs*^*14*^ harbored significantly more *E. faecalis* compared to the wild-type *w* (**Fig 3B**, p-adjusted = 0.032), and indeed *ΔAMPs*^*14*^ had bacterial abundances equivalent or even greater than *Rel*^*E20*^ flies for all bacterial taxa except *Le. pseudomesenteroides* (**Fig 3B**).

Overall, our data indicate that in the absence of bacterial community dynamics, AMPs and to a lesser extent lysozymes, are major effectors regulating gut microbiota abundance.

### Systemic infection with microbiota members

Previously, we showed that a lack of AMPs in the gut significantly affects the microbiota composition and growth. However, it is unclear whether AMPs have preferential antimicrobial activity that selects for core microbiota members, and to date it has not been demonstrated that AMPs directly control members of the microbiota community. In *Drosophila* AMPs are primarily active against Gram-negative bacteria, but less so against Gram-positive bacteria. However, it is not known whether AMPs can control bacterial species commonly found in the gut, with the exception of *E. faecalis* (45).

To address this, we used a systemic infection model to effectively “incubate” gut microbiota members in hemolymph with or without AMPs. Flies that fail to control bacterial proliferation ultimately die (66). We systemically infected flies with three representative bacteria that are normally present in the digestive tract and followed fly survival. We challenged wild-type, *ΔAMP*^*14*^*s, Rel*^*E20*^ and *spz*^*rm7*^ female flies by clean injury, and with three different bacterial species: *Acetobacter sp*. and *La. plantarum*, which have DAP-type peptidoglycan, and *E. faecalis* which has Lys-type peptidoglycan. *Rel*^*E20*^ lack a functional IMD response and are known to be very susceptible to systemic infection by most Gram-negative bacteria and certain classes of Gram-positive bacteria, while *spätzle* (*spz*^*rm7*^) mutants lack Toll immune signaling and are susceptible to Gram-positive bacteria and fungi. We observed that *ΔAMPs*^*14*^ flies were more susceptible to Gram-negative *Acetobacter sp*., mimicking the susceptibility of *Relish* mutants (**Fig 4**). As expected, *spz*^*rm7*^ flies were highly susceptible to Gram-positive *La. plantarum* and *E. faecalis* infection, dying completely within one week. However, *ΔAMPs*^*14*^ flies did not have increased mortality when infected with these bacterial species. Flies did not die upon clean injury, indicating that the phenotype is specific to bacterial infection and is not due to a technical bias in the experiment (**Fig S3**).

**Fig 4.**
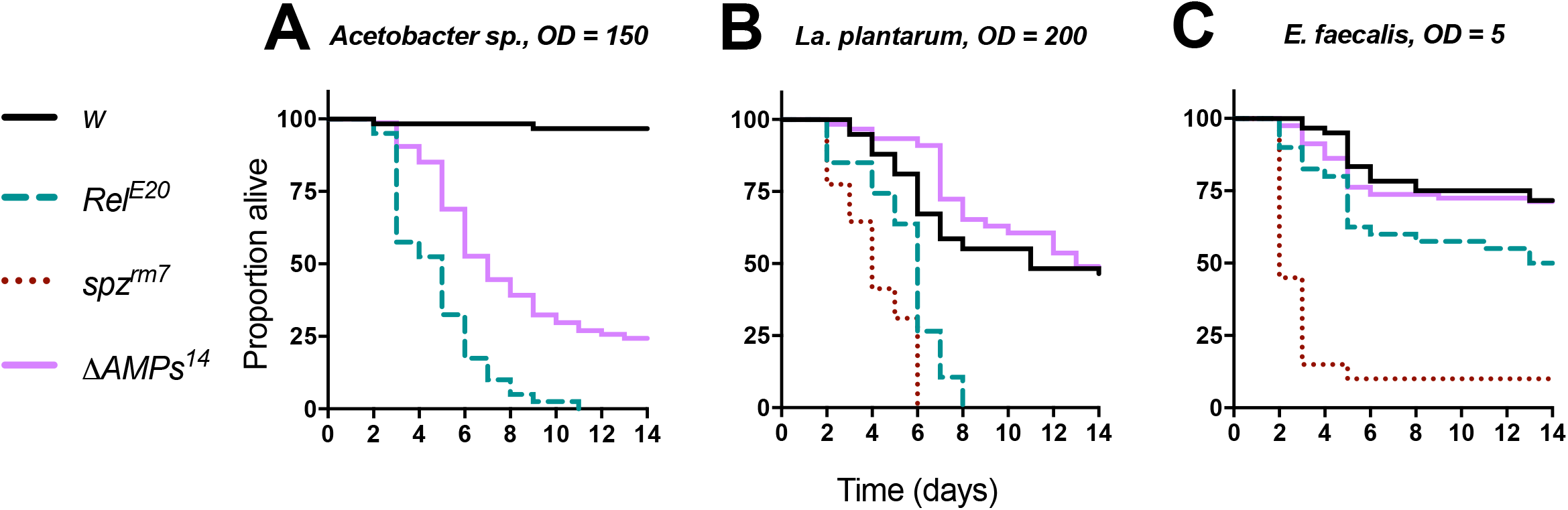
Survival upon systemic infection with microbiota bacteria. Female iso *w*^*1118*^ wild-type flies (*w*), and *Relish* (*Rel*^*E20*^), antimicrobial peptide (*ΔAMPs*^*14*^) and *spaetzle* (*spz*^*rm7*^) mutants were pricked in the thorax with three common microbiota bacteria: Gram-negative bacterium *Acetobacter sp*. (A) and two Gram-positive bacteria *La. Plantarum* (B) and *E. faecalis* (C). *ΔAMPs*^*14*^ mutants were significantly more susceptible than wild-type only to *Acetobacter sp*. infection (p = .001), and otherwise resisted infection like wild-type (p > 0.1). Pellet densities are reported for all systemic infections as OD600nm.

These systemic infections confirm that AMPs can play a direct role in the control of *Acetobacter sp*., bacteria typically found in the gut, but have a lesser impact on *La. plantarum* and *E. faecalis* proliferation. This trend is consistent with the results above showing that AMPs most prominently contribute to *Acetobacter* control after gnotobiotic or mono-associative colonization.

## Discussion

In *Drosophila*, the immune system and particularly the IMD pathway has been robustly demonstrated to be an important regulator of the gut microbiota and intestinal homeostasis (21, 22, 28, 30, 33, 37). Several reports have indicated the importance of the IMD pathway in maintaining balanced microbiota during aging, and mutants for this pathway (e.g *Relish*) have atypical microbiota abundance and composition (28). While it is clear that the IMD pathway is a regulator of the gut microbiota, little is known about the effectors mediating this regulation. In addition to regulating most AMP expression in the gut (21), the IMD pathway regulates other physiological aspects, including expression of digestive enzymes (42), and enterocyte delamination (40, 67, 68). The present work extends these studies by more narrowly defining the role of AMPs and lysozymes by comparing specific loss of these effectors to total loss of IMD signaling.

Our results confirm a prominent role for the IMD pathway in regulating both microbiota load and diversity, especially upon aging (21, 28, 37). The *ΔAMPs*^*14*^ mutant mimics the *Relish* phenotype in many respects, showing that AMPs indeed contribute downstream of IMD to shape the microbiota composition. Our experiments consistently showed an increase in the load of *Acetobacter* species in both *Rel*^*E20*^ and *ΔAMPs*^*14*^ flies. The observation that both *ΔAMPs*^*14*^ and *Rel*^*E20*^ flies are also susceptible to *Acetobacter* systemic infection, together with previous studies showing that AMPs contribute to survival to Gram-negative bacterial infection (45), provides strong evidence that direct microbicidal activity of AMPs regulates these Gram-negative bacteria. Collectively, this indicates for the first time that the basal level of IMD pathway activity induced by the gut microbiota (28) leads to the production of AMPs that prevent overgrowth of Gram-negative commensals such as *Acetobacter*. Future studies should clarify which AMP(s) among the 14 deleted in the *ΔAMPs*^*14*^ flies regulate *Acetobacter*.

*La. plantarum* is an important member of *Drosophila* microbiota that is associated to the host both in larval development and adulthood (11, 34, 69). As the DAP-type peptidoglycan found in the cell wall of these bacteria can activate the IMD pathway, we might expect to see an action of AMPs against them. However, the IMD pathway and AMP mutants are not very susceptible to DAP-type Gram-positive bacteria (70). Moreover, D-Alanylation of *La. plantarum* lipoteichoic acid has recently been proposed as a mechanism to protect against the action of AMPs and lysozymes (49). Here, we found that the AMPs play a role in controlling *La. plantarum* abundance in a mono-colonization setup but not when bacteria are in a community context. This might be due to the dynamics between microbiota members (e.g. competition between different species), or due to differential affinity of AMPs to the peptidoglycan of distinct species. It is possible that the abundance of *La. plantarum* is maintained at a threshold level in the gut and this is naturally achieved in a community through bacterial interactions. However, *La. plantarum* overgrowth can be inhibited by AMPs in a context where it becomes the only dominant member of the community.

The genome of *Drosophila* contains many genes encoding lysozymes, likely as a consequence of living in bacterially enriched habitats (46). Indeed, animals feeding on fermenting medium, such as ruminants or fruit flies, have a much higher number of lysozyme gene copies compared to animals feeding on ‘clean food’ (71, 72). In many insects, lysozymes are induced upon systemic infection, pointing to a possible role as immune effectors. In contrast, *Drosophila* lysozymes are strongly expressed in the gut, indicating a specific role in the digestive process (46, 47). Of note, one uncharacterized gene annotated as encoding a putative lysozyme (*CG6429*) is strongly induced upon systemic infection, and is partially regulated by the IMD pathway (73).

In this study, we generated a *LysB-P*^*Δ*^ mutant deficient for four lysozyme genes strongly expressed in the gut. *LysB-P*^*Δ*^ gut extracts have reduced lysozyme activity (**Fig S1**) confirming that these four genes indeed contribute to gut lysozyme activity. As lysozymes are known to digest peptidoglycan and can exhibit bactericidal activity alone or in combination with AMPs (48), we were interested to monitor the impact of lysozymes on the gut microbiota. We expected that loss of lysozymes would have a greater effect on Gram-positive bacteria, as the thin peptidoglycan layer of Gram-negative bacteria is protected by their external LPS membrane. Consistently, 16S sequencing revealed that *LysB-P*^*Δ*^ mutants exhibit increased relative community *Lactobacillus* abundance. However, mono-association experiments revealed a role of lysozymes in suppressing growth of only Gram-negative *Acetobacter* species. This effect was less marked than that of AMP deficient mutants.

An interesting observation of our study is that flies lacking AMPs or lysozymes display greater community stochasticity, similar to the phenotype of *Rel*^*E20*^ flies. This suggests that multiple factors including AMPs, lysozymes and bacteria-bacteria interactions contribute to stability of the gut microbiota, and that loss of these factors increases stochasticity. It should be noted that our work relies on the use of isogenic fly strains. While the isogenization process homogenizes the genetic background, it also increases the degree of homozygosity along the genome with a possible increase in genetic interactions. Thus, our study on AMPs and lysozymes using the *iso Drosdel* background should be reinforced by other studies using other backgrounds or alternative approaches.

In *Drosophila*, the induction of antibacterial peptides genes after infection is blocked in IMD pathway mutants, such as *Relish*, resulting in high susceptibility. These flies also cannot control their microbiota load, especially during aging (28). As expected, we found similar gut microbiota structure in *Relish* and *AMP* mutants. Indeed, both genotypes were unable to control the microbiota load and composition, but *Relish* flies had a more severe phenotype, with 16S analysis showing atypical microbial composition at early life stages, and marked inability to control all inoculated bacterial species in mono-association experiments. This is likely due to the multiple roles of the IMD pathway in gut physiology, apoptosis, nutrition and metabolism (40, 74, 75), the loss of which in addition to AMPs may exacerbate gut dysbiosis or hasten the inability of the flies to control microbiota growth. This indicates that although AMPs play an important role in control of microbiota members, they contribute only partially to the dysbiosis of IMD pathway mutant flies.

Collectively, our work is the first to show direct involvement of AMPs and lysozymes in the control of *Drosophila* gut microbiota. Consequences of the loss of these effectors are exacerbated during aging, and their loss contributes to increased microbiota abundance and shifted composition. This work shows that immune effectors typically associated with resistance to pathogenic infections also help shape the beneficial gut community, consistent with the idea that host-symbiont interactions use the same ‘language’ typically associated with pathogenesis (76).

## Material and Methods

### Bacterial strains and culture conditions

Bacterial strains used in this study and their origins are as follows: *Acetobacter sp*. (52), *Acetobacter pomorum* (50), *Lactobacillus plantarum* (11), *Enterococcus faecalis* (51). *Lactobacillus brevis* and *Leuconostoc pseudomesenteroides* were isolated from the “Valais” population, collected in the Valais canton of Switzerland in 2007 (77). Briefly, homogenates from 20 flies were spread over MRS-5% Mannitol plates. A single colony was used to prepare liquid cultures (described below) and establish glycerol stocks, as well as for 16S rRNA gene full-length amplification using universal primers. The PCR products were sequenced by Sanger sequencing and assigned to taxa based on a Microbial BLASTn search against the nucleotide database of NCBI (https://blast.ncbi.nlm.nih.gov/Blast.cgi?PAGE_TYPE=BlastSearch&BLAST_SPEC=MicrobialGenomes). See **Supplementary Text 1** for full 16S rRNA gene sequences of these isolates and *Acetobacter* sp (52).

*A. pomorum* and *Acetobacter sp*. were cultivated in Man, Rogosa and Sharpe (MRS)-D-Mannitol 2,5% medium in aerobic conditions at 29°C for at least 18h with agitation. *La. plantarum, La. brevis* and *Le. pseudomesenteroides* were cultivated in anaerobic conditions in (MRS)-D-Mannitol 2,5% medium at 29°C for at least 18h standing. *E. faecalis* was cultured in Brain Heart Infusion Broth (BHI) medium in aerobic conditions at 37°C for at least 16h with agitation.

### Gnotobiotic fly cultivation and media

All experiments were done on flies kept at 25°C with a 12h dark/light cycle.

Because antibiotic treatments over multiple generations may result in epigenetic effects that may interfere with phenotypes, GF flies were freshly generated for each experiment. Please see extended Material and Methods for details.

To avoid sticky ‘biofilm-like’ formation on the medium, flies were transferred to fresh medium every 3-4 days. To avoid a decrease in microbial loads, the microbiota of each tube was transferred to the next using glass beads. Briefly, flies were anaesthetized on ice and removed on sterile caps. 10-20 glass beads were transferred to the old tube and shaken for 10 seconds.

The beads were then transferred to the new sterile tube and were shaken again for 10 seconds to spread the bacteria around the tubes. Adults were added in the new tubes. Flies were sampled 10 days and 29 days after colonization.

For vertically transmitted microbiota, we let gnotobiotic adults (colonized as described above 3-4 days previously) lay eggs on fresh medium for three days. We let larvae grow in this original medium and began transferring the emerging adults to new tubes 5 days after the first fly emerged. We collected adults 10-15 days after emergence and analyzed their associated bacterial community.

For mono-association experiments, we colonized flies with each isolate that was included in the commensal cocktail. Because during mono-association changes in community structure is not a concern, we maintained the flies in their original tube throughout the experiment. Flies were sampled 6 days after colonization.

### DNA extraction and qPCR

DNA extraction was carried out on samples of surface sterilized adults (washed in sterile water and EtOH) using a DNeasy Blood & Tissue Kit (Qiagen).

The qPCR reactions for absolute quantification were carried out as described in (78) and for relative quantification qPCR reactions as described in (65). The universal (78) and *Acetobacter* specific 16S (65), and Actin 5C primers (65) were previously described. Firmicute specific primers (anti-sense primer 5’AGCGTTGTCCGGATTTAT 3’, sense primer 5’ CATTTCACCGCTACACAT 3’) were designed by aligning the 16S rRNA gene sequences of the four Firmicute species that were used in this study. Their specificity (lack of amplification in *Acetobacteraceae*) was determined on plasmid DNA containing specific 16S sequences as well as on DNA extracted from flies mono-associated with microbiota members described in this study.

### 16S rRNA gene amplicon sequencing and data processing

Amplification of the V4 region of the 16S rRNA gene and library preparation protocol was done as previously described (78) except for the first cycle PCR that was performed with 30 cycles. Libraries were verified by Fragment analyzer, mixed with 10% PhiX library (Illumina #FC-110-3001), and subjected to Illumina MiSeq v3 paired-end sequencing in one lane, with all libraries multiplexed.

Sequencing data has been processed using Divisive Amplicon Denoising Algorithm 2 (DADA2) pipeline (“dada2” package version 1.14.1 in R) and “phyloseq” package version 1.30.0. Please see extended methods for further details.

### Diversity and statistical analysis

Permutational multivariate analysis of variance (ADONIS, “adonis” function) based on Bray– Curtis distances (“vegdist” function) (79) was used to test the effects of age and genotype on community structure, and “metaMDS” function was used for plotting beta-diversity. For pairwise comparisons of ADONIS, “adonis.pair” function was used from “EcolUtils” package. To test the dispersion of communities we used the function “betadisper” (80, 81) and compared the distances of individual samples to group centroids in multidimensional space using “permutest”.

All statistical analyses were performed using R (version 3.6.3). We used general linear mixed models (“lme4” package version 1.1.23) to test for the effects of age, genotype and their interaction (depending on the experimental design) on bacterial loads or dispersion of bacterial communities. Pairwise comparisons were performed using “emmeans” and “pairs” functions (“emmeans” package version 1.5.1). p-values were adjusted using FDR method.

### Systemic infection and lifespan assay

Systemic infections were performed by pricking 5-7 day-old conventionally reared adult females in the thorax with a 100-μm-thick insect pin dipped into a concentrated pellet of bacteria. The following bacteria were grown to the respective OD600 concentrations in (MRS)-D-Mannitol 2,5% broth (*Acetobacter sp*., OD 600=150; *La. plantarum*, OD600=200) or BHI (*E*.*faecalis*, OD600=5). Infected flies were maintained at 25°C for experiments. At least three replicate survival experiments were performed for each infection, with 20 flies per vial on standard fly medium without yeast. Survivals were scored daily and flies were flipped to fresh medium every two days.

Survival data were analyzed using the survival package in R 3.6.3 with a Cox proportional hazards model (coxph() function) including experiment as a covariate in the final model when significant.

## Acknowledgments

We are grateful to Tadeusz J. Kawecki (University of Lausanne, Switzerland) for providing bacterial isolates; Xiaoxue Li for providing data on *LysB-P*^*Δ*^ strain validation; Iatsenko Igor (Max Planck Institute, Germany) for *LysB-P*^*Δ*^ strain isogenization; and to Hannah Westlake for critical reading of the manuscript.

This work was supported by the Swiss National Science Foundation grant N°310030_185295. The funders had no role in study design, data collection and interpretation, or the decision to submit the work for publication

## Author contributions

AM, BE, BL conceived the study and designed the experiments; AM, BE, MH, collected the data; BE analyzed the data; SK provided critical reagents; AM, BE, MH and BL wrote the manuscript.

## Data availability

16S rRNA gene amplicon sequencing data will be submitted to NCBI database upon acceptance.

## Figure legends

**Fig Supp1. Lysozyme mutant characterization**.

A) Diagram of the *LysB-P*^*Δ*^ mutation, which deletes 11.5kb of nucleotide sequence from the *Drosophila* genome. *LysB-P*^*Δ*^ flies lack the four lysozyme genes (*LyB, LysD, LysE, LysP*) that are highly expressed in the gut, and also the *LysC* pseudogene. B) *LysB-P*^*Δ*^ mutants have a reduced capacity to degrade peptidoglycan in the gut compared to wild-type flies. Gut extracts from wild-type flies (*w*) and lysozyme mutants (*LysB-P*^*Δ*^) were incubated with *E*.*faecalis* peptidoglycan for 48h. The lysozyme activity was monitored by change in optical density of the peptidoglycan solution. Commercial lysozyme was included as a positive control.

**Fig Supp2. Homogeneity of multivariate dispersals in microbial communities**.

Distance to the center of each group (genotype) represents how spread are the communities in the multivariate space (PCoA) in A) a vertical transmission setup, B) 10 days after colonization, and C) 29 days after colonization. Different letters indicate significant differences (p<0.05) between p-values upon pairwise comparison, calculated by permutations on multivariate equivalent of variances. A significant difference between groups indicates that samples belonging to one group are more variable in microbiota composition compared to the other group.

**Fig Supp3. *ΔAMPs***^***14***^ **flies survive clean injury**.

Females from indicated genotypes were pricked with a clean 100μm insect pin. No genotype displayed pronounced susceptibility to clean injury.

**Supplementary Text 1:** 16S rRNA gene sequences of *La. brevis, Le. Pseudomesenteroides* and *Acetobacter sp*. [REF erkosar 2017] used in this study.

**Supplementary Table S1**: number of flies used in each samples and number of reads that are retained after each step of 16S rRNA gene analysis

**Supplementary Table S2**: Summary of the statistics used for the data analysed in Fig1. and Fig2.

