## Supplementary figures and images for "*Drosophila* antimicrobial peptides and lysozymes regulate gut microbiota composition and abundance"

### Fig Supp1.

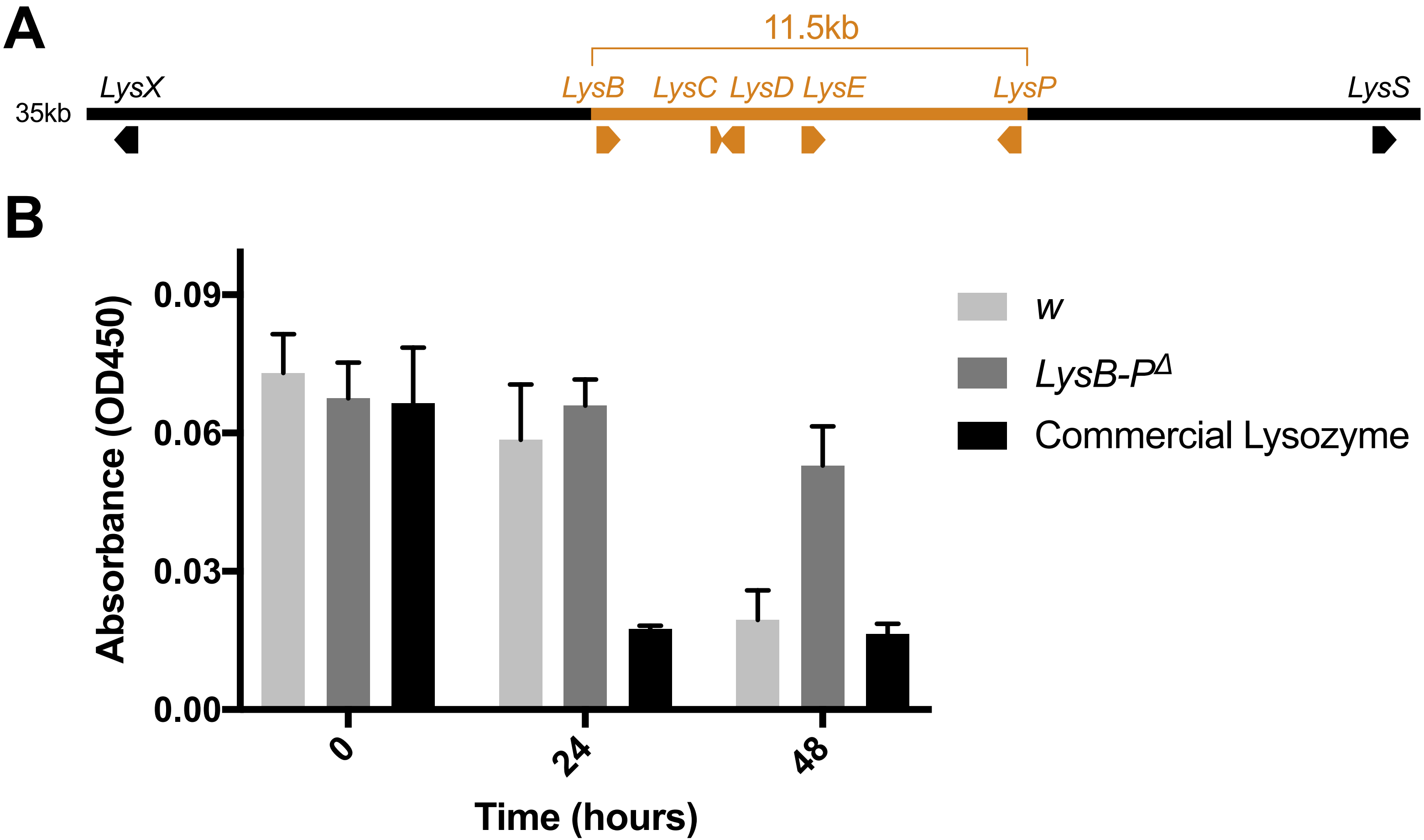

### Fig Supp2.

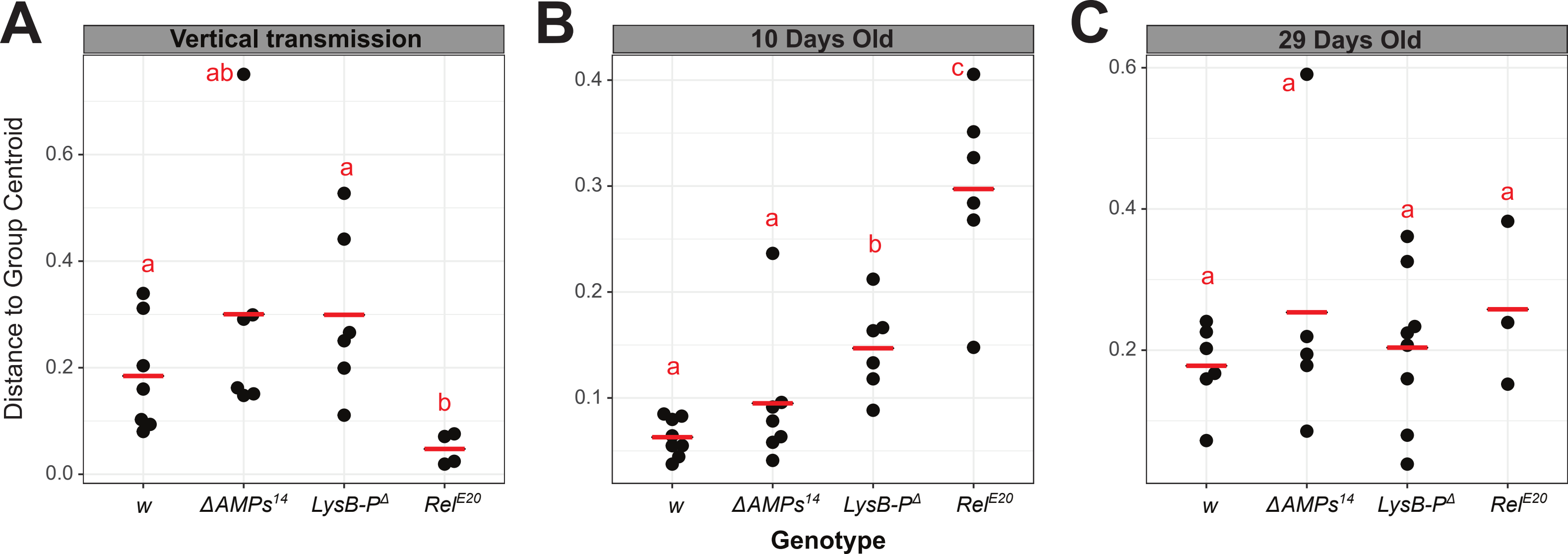

### Fig Supp3.

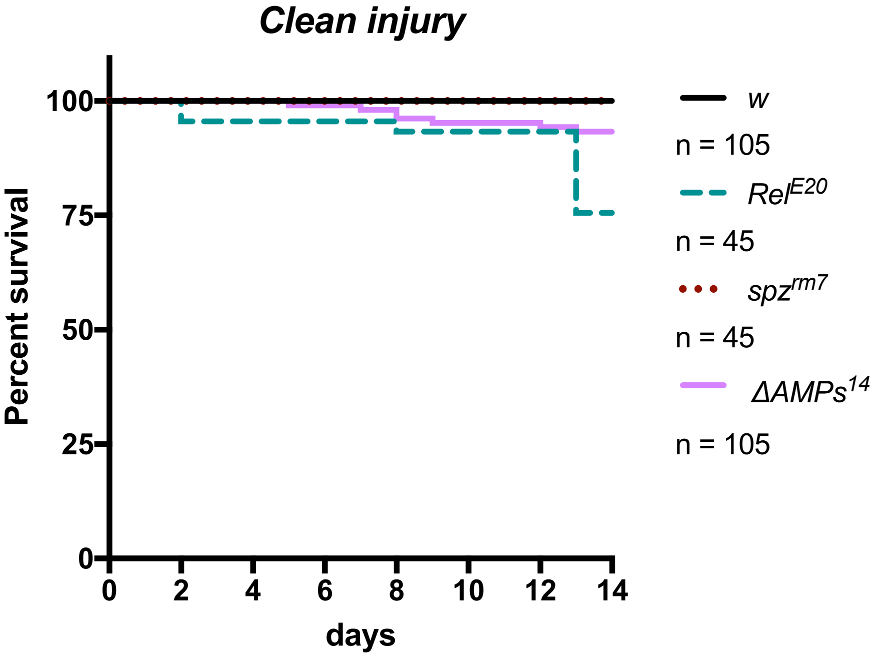
