## Supplementary Text 1 for "*Drosophila* antimicrobial peptides and lysozymes regulate gut microbiota composition and abundance"

**Supplementary Text 1:** 16S rRNA gene sequences of La. brevis, *Le. pseudomesenteroides* and *Acetobacter sp*. [REF erkosar 2017] used in this study.

>Lactobacillus_brevis_16S
CCTAATACATGCAAGTCGAACGAGCTTCCGTTGAATGACGTGCTTGCACTGATTTCAACAATGAAGC
GAGTGGCGAACTGGTGAGTAACACGTGGGGAATCTGCCCAGAAGCAGGGGATAACACTTGGAAAC
AGGTGCTAATACCGTATAACAACAAAATCCGCATGGATTTTGTTTGAAAGGTGGCTTCGGCTATCAC
TTCTGGATGATCCCGCGGCGTATTAGTTAGTTGGTGAGGTAAAGGCCCACCAAGACGATGATACGT
AGCCGACCTGAGAGGGTAATCGGCCACATTGGGACTGAGACACGGCCCAAACTCCTACGGGAGGC
AGCAGTAGGGAATCTTCCACAATGGACGAAAGTCTGATGGAGCAATGCCGCGTGAGTGAAGAAGG
GTTTCGGCTCGTAAAACTCTGTTGTTAAAGAAGAACACCTTTGAGAGTAACTGTTCAAGGGTTGACG
GTATTTAACCAGAAAGCCACGGCTAACTACGTGCCAGCAGCCGCGGTAATACGTAGGTGGCAAGCG
TTGTCCGGATTTATTGGGCGTAAAGCGAGCGCAGGCGGTTTTTTAAGTCTGATGTGAAAGCCTTCGG
CTTAACCGGAGAAGTGCATCGGAAACTGGGAGACTTGAGTGCAGAAGAGGACAGTGGAACTCCAT
GTGTAGCGGTGGAATGCGTAGATATATGGAAGAACACCAGTGGCGAAGGCGGCTGTCTAGTCTGT
AACTGACGCTGAGGCTCGAAAGCATGGGTAGCGAACAGGATTAGATACCCTGGTAGTCCATGCCGT
AAACGATGAGTGCTAAGTGTTGGAGGGTTTCCGCCCTTCAGTGCTGCAGCTAACGCATTAAGCACTC
CGCCTGGGGAGTACGACCGCAAGGTTGAAACTCAAAGGAATTGACGGGGGCCCGCACAAGCGGTG
GAGCATGTGGTTTAATTCGAAGCTACGCGAAGAACCTTACCAGGTCTTGACATCTTCTGCCAATCTTA
GAGATAAGACGTTCCCTTCGGGGACAGAATGACAGGTGGTGCATGGTTGTCGTCAGCTCGTGTCGT
GAGATGTTGGGTTAAGTCCCGCAACGAGCGCAACCCTTATTATCAGTTGCCAGCATTCAGTTGGGCA
CTCTGGTGAGACTGCCGGTGACAAACCGGAGGAAGGTGGGGATGACGTCAAATCATCATGCCCCTT
ATGACCTGGGCTACACACGTGCTACAATGGACGGTACAACGAGTTGCGAAGTCGTGAGGCTAAGCT
AATCTCTTAAAGCCGTTCTCAGTTCGGATTGTAGGCTGCAACTCGCCTACATGAAGTTGGAATCGCTA
GTAATCGCGGATCAGCATGCCGCGGTGAATACGTTCCCGGGCCTTGTACACACCGCCCGTCACACCA
TGAGAGTTTGTAACACCCAAAGCCGGTGAGATAACCTTCGGGAGTCAGCCGTCTAAGGGACGAT

>Leuconostoc_pseudomesenteroides_16S
GCTAATACATGCAAGTCGAACGCACAGCGAAAGGTGCTTGCACCTTTCAGTGAGTGGCGAACGGGT
GAGTAACACGTGGACAACCTGCCTCAAGGCTGGGGATAACATTTGGAAACAGATGCTAATACCGAA
TAAAACTCAGTGTCGCATGACACAAAGTTAAAAGGCGCTTTGGCGTCACCTAGAGATGGATCCGCG
GTGCATTAGTTAGTTGGTGGGGTAAAGGCCTACCAAGACAATGATGCATAGCCGAGTTGAGAGACT
GATCGGCCACATTGGGACTGAGACACGGCCCAAACTCCTACGGGAGGCTGCAGTAGGGAATCTTCC
ACAATGGGCGAAAGCCTGATGGAGCAACGCCGCGTGTGTGATGAAGGCTTTCGGGTCGTAAAGCA
CTGTTGTATGGGAAGAACAGCTAGAATAGGGAATGATTTTAGTTTGACGGTACCATACCAGAAAGG
GACGGCTAAATACGTGCCAGCAGCCGCGGTAATACGTATGTCCCGAGCGTTATCCGGATTTATTGG
GCGTAAAGCGAGCGCAGACGGTTGATTAAGTCTGATGTGAAAGCCCGGAGCTCAACTCCGGAATG
GCATTGGAAACTGGTTAACTTGAGTGCAGTAGAGGTAAGTGGAACTCCATGTGTAGCGGTGGAATG
CGTAGATATATGGAAGAACACCAGTGGCGAAGGCGGCTTACTGGACTGTAACTGACGTTGAGGCTC
GAAAGTGTGGGTAGCAAACAGGATTAGATACCCTGGTAGTCCACACCGTAAACGATGAACACTAGG
TGGTTAGGAGGTTTCCGCCTCTTAGTGCCGAAGCTAACGCATTAAGTGTTCCGCCTGGGGAGTACGA
CCGCAAGGTTGAAACTCAAAGGAATTGACGGGGACCCGCACAAGCGGTGGAGCATGTGGTTTAATT
CGAAGCAACGCGAAGAACCTTACCAGGTCTTGACATCCTTTGAAGCTTTTAGAGATAGAAGTGTTCT
CTTCGGAGACAAAGTGACAGGTGGTGCATGGTCGTCGTCAGCTCGTGTCGTGAGATGTTGGGTTAA
GTCCCGCAACGAGCGCAACCCTTATTGTTAGTTGCCAGCATTCAGATGGGCACTCTAGCGAGACTGC
CGGTGACAAACCGGAGGAAGGCGGGGACGACGTCAGATCATCATGCCCCTTATGACCTGGGCTACA
CACGTGCTACAATGGCGTATACAACGAGTTGCCAACCCGCGAGGGTGAGCTAATCTCTTAAAGTAC

GTCTCAGTTCGGATTGTAGTCTGCAACTCGACTACATGAAGTCGGAATCGCTAGTAATCGCGGATCA
GCACGCCGCGGTGAATACGTTCCCGGGTCTTGTACACACCGCCCGTCACACCATGGGAGTTTGTAAT
GCCCAAAGCCGGTGGCCTAACCTTTAGGAAGGAGCCGTCTAAGGCAGACCG

>Acetobacter_sp_16S
atGCaGTcgcanGaaGgCTTCGGCCTTAGTGGCGGacGGGTGAGTAACGCGTAGGAATCTATCCatGGg
tGGGGGATAACTCCGGGAAACTGGAGctAATACCGCATGATACCTGAGGGTCAAAGGCGCAAGTCG
CCTgtGGAGGAGCCTGCGTTTGATTAGCTTGTTGGTGGGGTAAAGGCCTACCAAGGCGATGATCAAT
AGCTGGTCTGAGAGGATGATCAGCCACACTGGGACTGAGACACGGCCCAGACTCCTACGGGAGGC
AGCAGTGGGGAATATTGGACAATGGGGGAAACCCTGATCCAGCAATGCCGCGTGTGTGAAGAAGG
TTTTCGGATTGTAAAGCACTTTCGGCGGGGACGATGATGACGGTACCCGCAGAAGAAGCCCCGGCT
AACTTCGTGCCAGCAGCCGCGGTAATACGAAGGGGGCTAGCGTTGCTCGGAATGACTGGGCGTAA
AGGGCGTGTAGGCGGTTTGTACAGTCAGATGTGAAATCCCCGGGCTTAACCTGGGAGCTGCATTTG
ATACGTGCAGACTAGAGTATGAGAGAGGGTTGTGGAATTCTCAGTGTAGAGGTGAAATTCGTAGAT
ATTGGGAAGAACACCGGTGGCGAAGGCGGCAACCTGGCTCATTACTGACGCTGAGGCGCGAAAGC
GTGGGGAGCAAACAGGATTAGATACCCTGGTAGTCCACGCTGTAAACGATGTGTGCTGGATGTTGG
GTAACTTAGTTACTCAGTGTCGTAGCTAACGCGATAAGCACACCGCCTGGGGAGTACGGCCGCAAG
GTTGAACTCAAAGGAATTGACGGGGGCCCGCACAAGCGGTGGAGCATGTGGTTTAATTCGAAGCA
ACGCGCAGAACCTTACCAGGGCTTGTATGGAGAGGCTGTATTCAGAGATGGATATTTCCCGCAAGG
GACCTCTTGCACAGGTGCTGCATGGCTGTCGTCAGCTCGTGTCGTGAGATGTTGGGTTAAGTCCCGC
AACGAGCGCAACCCTTATCTTTAGTTGCCAGCATGTTTGGGTGGGCACTCTAAAGAGACTGCCGGTG
ACAAGCCGGAGGAAGGTGGGGATGACGTCAAGTCCTCATGGCCCTTATGTCCTGGGCTACACACGT
GCTACAATGGCGGTGAcagTGGGAAGCTAGATGGTGACATCATGCCGATCTCTAAAAACCGTCTCAG
TTCGGATTGCACTCTGCAACTCGAGTGCATGAAGGTGGAATCGCTAGTAATCGCGGATCAGcatGCC
GCGGTGAATACGTTCCCGGGCCTTGTACAcacCGCCCGTCACaccATGGGAGTTGGnTnaCCTTAAGC
CgGTgaGCGAACCgcaaGAcgcaAGcgGa
