## Supplementary Table S1 for "*Drosophila* antimicrobial peptides and lysozymes regulate gut microbiota composition and abundance"

| Sample* | # Flies | initial #reads | Filtered | Denoised F | Denoised R | Merged | Non-chimeras | bacFiltered** |
|---|---|---|---|---|---|---|---|---|
| dAMP_1_r1_10d_1 | 10 | 198415 | 119889 | 119831 | 119589 | 117941 | 117941 | 117231 |
| dAMP_1_r1_10d_2 | 10 | 279472 | 165273 | 165226 | 164800 | 163304 | 163304 | 163040 |
| dAMP_1_r1_29d_1 | 8 | 136085 | 81191 | 81051 | 80971 | 79325 | 77216 | 77168 |
| dAMP_1_r1_VerticalTrans_1 | 5 | 134880 | 82475 | 82440 | 82332 | 82003 | 82003 | 81882 |
| dAMP_1_r2_10d_1 | 10 | 221248 | 134917 | 134865 | 134711 | 133002 | 133002 | 132683 |
| dAMP_1_r2_10d_2 | 5 | 285526 | 174552 | 174449 | 174191 | 171816 | 170349 | 167301 |
| dAMP_1_r2_29d_1 | 6 | 270998 | 164534 | 164494 | 164341 | 159733 | 159502 | 159425 |
| dAMP_1_r2_29d_2 | 6 | 216330 | 131327 | 131261 | 131102 | 126716 | 126361 | 126312 |
| dAMP_1_r2_VerticalTrans_1 | 7 | 150429 | 92007 | 91961 | 91771 | 91457 | 91266 | 91232 |
| dAMP_1_r3_VerticalTrans_1 | 8 | 123405 | 74463 | 74421 | 74275 | 73952 | 73952 | 73811 |
| dAMP_2_r1_10d_1 | 10 | 239915 | 144406 | 144346 | 144124 | 143049 | 142999 | 140812 |
| dAMP_2_r1_10d_2 | 4 | 314766 | 187147 | 187010 | 186723 | 184739 | 184572 | 175126 |
| dAMP_2_r1_29d_1 | 7 | 344596 | 206625 | 205146 | 206151 | 197609 | 191977 | 191559 |
| dAMP_2_r1_VerticalTrans_1 | 6 | 164042 | 99782 | 99734 | 99599 | 98849 | 98175 | 98169 |
| dAMP_2_r2_10d_1 | 9 | 140966 | 85716 | 85678 | 85600 | 84943 | 84943 | 84596 |
| dAMP_2_r2_29d_1 | 10 | 122801 | 73717 | 73636 | 73468 | 71662 | 70645 | 70578 |
| dAMP_2_r2_VerticalTrans_1 | 10 | 170047 | 99629 | 99524 | 99231 | 97980 | 96977 | 96824 |
| dAMP_2_r3_VerticalTrans_1 | 8 | 333021 | 200848 | 200737 | 200405 | 199301 | 197995 | 197792 |
| Lys_1_r1_10d_1 | 10 | 288237 | 176047 | 175950 | 175743 | 170804 | 170376 | 168492 |
| Lys_1_r1_10d_2 | 10 | 288783 | 176693 | 176652 | 176449 | 171898 | 171584 | 170195 |
| Lys_1_r1_29d_1 | 10 | 215621 | 127915 | 127857 | 127617 | 122928 | 122911 | 122670 |
| Lys_1_r1_29d_2 | 8 | 139254 | 83476 | 83390 | 83228 | 79880 | 79875 | 79591 |
| Lys_1_r1_VerticalTrans_1 | 8 | 278798 | 161179 | 161141 | 160870 | 158477 | 158274 | 152621 |
| Lys_1_r2_10d_1 | 10 | 213270 | 129430 | 129333 | 129179 | 125752 | 125462 | 124567 |
| Lys_1_r2_10d_2 | 10 | 386433 | 231518 | 231443 | 231077 | 222268 | 221365 | 220796 |
| Lys_1_r2_29d_1 | 10 | 146137 | 87704 | 87647 | 87447 | 84829 | 84408 | 84183 |
| Lys_1_r2_29d_2 | 8 | 200615 | 121173 | 121071 | 120898 | 117296 | 117248 | 116867 |

| | | | | | | | | |
|---|---|---|---|---|---|---|---|---|
| Lys_1_r2_VerticalTrans_1 | 4 | 283688 | 168556 | 168487 | 167839 | 165812 | 165645 | 165410 |
| Lys_1_r3_VerticalTrans_1 | 5 | 334811 | 196729 | 196645 | 196016 | 193442 | 193240 | 192974 |
| Lys_2_r1_10d_1 | 10 | 166350 | 100089 | 100023 | 99741 | 98802 | 98792 | 98307 |
| Lys_2_r1_10d_2 | 10 | 102189 | 60615 | 60566 | 60449 | 59555 | 59555 | 59344 |
| Lys_2_r1_29d_1 | 10 | 221088 | 134220 | 134163 | 133938 | 130293 | 130219 | 130054 |
| Lys_2_r1_29d_2 | 10 | 160493 | 96178 | 96082 | 95819 | 93027 | 92910 | 92727 |
| Lys_2_r1_VerticalTrans_1 | 9 | 220361 | 133215 | 133191 | 132913 | 128959 | 128787 | 128745 |
| Lys_2_r2_29d_1 | 8 | 207383 | 124019 | 123937 | 123695 | 119557 | 118835 | 118659 |
| Lys_2_r2_29d_2 | 8 | 124609 | 75638 | 75582 | 75440 | 72762 | 72746 | 72657 |
| Lys_2_r2_VerticalTrans_1 | 10 | 231503 | 140724 | 140623 | 140538 | 137356 | 137356 | 136542 |
| Lys_2_r3_VerticalTrans_1 | 10 | 237096 | 142105 | 142036 | 141799 | 138235 | 138135 | 137701 |
| Rel_1_r1_10d_1 | 10 | 165672 | 99828 | 99776 | 99601 | 96875 | 96680 | 96365 |
| Rel_1_r1_29d_1 | 5 | 142222 | 83914 | 83858 | 83656 | 81733 | 77845 | 77832 |
| Rel_1_r2_10d_1 | 10 | 177025 | 105763 | 105706 | 105425 | 103224 | 103183 | 103065 |
| Rel_1_r2_10d_2 | 5 | 129464 | 74479 | 74409 | 74147 | 73186 | 72865 | 72677 |
| Rel_1_r2_VerticalTrans_1 | 8 | 180773 | 111286 | 111227 | 111085 | 110664 | 110664 | 110476 |
| Rel_1_r3_VerticalTrans_1 | 10 | 241828 | 148237 | 148134 | 147949 | 146704 | 146588 | 142552 |
| Rel_2_r1_10d_1 | 9 | 81990 | 45776 | 45756 | 45581 | 44972 | 44972 | 26135 |
| Rel_2_r1_10d_2 | 9 | 186257 | 108730 | 108680 | 108277 | 107542 | 107542 | 105403 |
| Rel_2_r1_29d_1 | 6 | 113524 | 68345 | 68304 | 68138 | 66029 | 62578 | 62576 |
| Rel_2_r1_VerticalTrans_1 | 10 | 293351 | 177116 | 177037 | 176758 | 175394 | 175277 | 173631 |
| Rel_2_r2_10d_1 | 10 | 287915 | 175016 | 174980 | 174737 | 173714 | 173670 | 170228 |
| Rel_2_r2_29d_1 | 2 | 92415 | 54838 | 54664 | 54474 | 51964 | 45982 | 42596 |
| Rel_2_r3_VerticalTrans_1 | 10 | 200504 | 120738 | 120666 | 120475 | 119917 | 119756 | 119667 |
| w_1_r1_10d_1 | 10 | 128717 | 78558 | 78511 | 78429 | 76910 | 76910 | 76720 |
| w_1_r1_10d_2 | 10 | 143826 | 87902 | 87853 | 87687 | 86171 | 86171 | 84858 |
| w_1_r1_29d_1 | 6 | 207126 | 125673 | 125594 | 125490 | 121147 | 121147 | 120730 |
| w_1_r1_29d_2 | 6 | 363182 | 219169 | 219066 | 218786 | 210104 | 210021 | 209582 |

| | | | | | | | | |
|---|---|---|---|---|---|---|---|---|
| w_1_r1_VerticalTrans_1 | 5 | 266497 | 156547 | 156497 | 155853 | 153045 | 152729 | 148294 |
| w_1_r2_10d_1 | 9 | 105442 | 63734 | 63650 | 63589 | 62063 | 61933 | 61359 |
| w_1_r2_10d_2 | 9 | 301687 | 184604 | 184554 | 184345 | 180408 | 180178 | 174903 |
| w_1_r2_29d_1 | 10 | 120509 | 72680 | 72612 | 72536 | 70235 | 70074 | 69904 |
| w_1_r2_29d_2 | 10 | 160109 | 97560 | 97493 | 97357 | 94195 | 93970 | 93833 |
| w_1_r2_VerticalTrans_1 | 4 | 206267 | 120736 | 120624 | 120222 | 118275 | 118124 | 117301 |
| w_1_r3_VerticalTrans_1 | 10 | 398820 | 232817 | 232734 | 231946 | 230352 | 227504 | 226582 |
| w_2_r1_10d_1 | 9 | 192644 | 115370 | 115332 | 115177 | 114343 | 114343 | 113117 |
| w_2_r1_10d_2 | 9 | 279236 | 168568 | 168517 | 168218 | 166890 | 166869 | 166488 |
| w_2_r1_29d_1 | 8 | 155685 | 95016 | 94908 | 94766 | 92547 | 92408 | 91946 |
| w_2_r1_VerticalTrans_1 | 10 | 258499 | 150454 | 150375 | 150009 | 147933 | 147770 | 144601 |
| w_2_r2_10d_1 | 10 | 248622 | 150758 | 150685 | 150480 | 149184 | 149184 | 147344 |
| w_2_r2_10d_2 | 4 | 378800 | 228769 | 228707 | 228452 | 226019 | 226014 | 224698 |
| w_2_r2_29d_1 | 10 | 157947 | 95484 | 95413 | 95275 | 91478 | 91478 | 91234 |
| w_2_r2_VerticalTrans_1 | 10 | 142551 | 86080 | 86013 | 85872 | 83170 | 83170 | 81939 |
| w_2_r3_VerticalTrans_1 | 10 | 412044 | 244344 | 244272 | 243626 | 237903 | 237026 | 236340 |
| w_2_r4_VerticalTrans_1 | 10 | 116841 | 68387 | 68299 | 68072 | 66661 | 66645 | 66111 |

* Genotype_Colonization Batch_Vial_Age/Life stage_sample replicate

** Reads that were matching tomitochondria, chloroplast or eukaryotes have been removed
