## Supplementary Table S2 for "*Drosophila* antimicrobial peptides and lysozymes regulate gut microbiota composition and abundance"

**MAIN INTERACTION MODEL**

**ADONIS on relative abundances with genotype and age as interacting factors, 999 Permutatio**

| | Df | SumsOfSqs | MeanSqs | F Model | R2 | P-value |
|---|---|---|---|---|---|---|
| GENOTYPE | 3 | 1.0456 | 0.34855 | 6.678 | 0.17489 | 0.002 |
| AGE | 1 | 2.3922 | 2.39216 | 45.835 | 0.40009 | 0.001 |
| GENOTYPE X AGE | 3 | 0.4014 | 0.13379 | 2.564 | 0.06713 | 0.03 |
| RESIDUALS | 41 | 2.1398 | 0.05219 | | 0.35789 | |

Pairwise ADONIS (different model) between ages, 1000 permutations

| | SumsOfSq | MeanSqs | F Model | R2 | P-value |
|---|---|---|---|---|---|
| 10d - 29d | 2.401378 | 2.401378 | 31.54751 | 0.401636 | 0.001 |

**10 DAYS OLD FLIES**

**ADONIS on relative abundances with genotype as a factor, 999 Permutations**

| | Df | F Model | R2 | P-value |
|---|---|---|---|---|
| GENOTYPE | 3 | 4.68 | 0.379 | 0.001 |
| RESIDUALS | 23 | | 0.621 | |

Pairwise ADONIS (different model) between genotypes, 1000 permutations

| | F Model | R2 | P-value | P-adjusted (FDR) |
|---|---|---|---|---|
| dAMP-Lys | 2.2833 | 0.17189 | 0.090909 | 0.109091 |
| dAMP-Rel | 4.2014 | 0.27638 | 0.042957 | 0.077922 |
| dAMP-w | 1.8614 | 0.12525 | 0.144855 | 0.144855 |
| Lys-Rel | 3.434 | 0.25562 | 0.051948 | 0.077922 |
| Lys-w | 5.3992 | 0.31031 | 0.002997 | 0.017982 |
| Rel-w | 7.8648 | 0.39592 | 0.005994 | 0.017982 |

**Homogeneity of multivariate dispersions with Genotype as a factor, 1000 permutations**

| | Df | F Model | P-value |
|---|---|---|---|
| GENOTYPE | 3 | 21.1 | 0.001 |
| RESIDUALS | 23 | | |

Pairwise comparisions from the same model:

| | Permuted P-value |
|---|---|
| dAMP-Lys | 0.125 |
| dAMP-Rel | 0.002 |
| dAMP-w | 0.230 |
| Lys-Rel | 0.003 |
| Lys-w | 0.000 |
| Rel-w | 0.000 |

**29 DAYS OLD FLIES**

**ADONIS on relative abundances with genotype as a factor, 999 Permutations**

| | Df | F Model | R2 | P-value |
|---|---|---|---|---|
| GENOTYPE | 3 | 4.26 | 0.415 | 0.008 |
| RESIDUALS | 18 | | 0.585 | |

Pairwise ADONIS (different model) between genotypes, 1000 permutations

| | F Model | R2 | P-value | P-adjusted (FDR) |
|---|---|---|---|---|
| dAMP-Lys | 1.7118 | 0.134662 | 0.195804 | 0.29371 |
| dAMP-Rel | 0.29448 | 0.046784 | 0.926074 | 0.92607 |
| dAMP-w | 8.3592 | 0.481543 | 0.01998 | 0.03996 |
| Lys-Rel | 1.04833 | 0.104329 | 0.402597 | 0.48312 |
| Lys-w | 7.72731 | 0.391706 | 0.013986 | 0.03996 |
| Rel-w | 9.40865 | 0.573396 | 0.018981 | 0.03996 |

**Homogeneity of multivariate dispersions with Genotype as a factor, 1000 permutations**

| | Df | F Model | P-value |
|---|---|---|---|
| GENOTYPE | 3 | 0.49 | 0.72 |
| RESIDUALS | 18 | | |

Pairwise comparisions from the same model:

| | Permuted P-value |
|---|---|
| dAMP-Lys | 0.567 |
| dAMP-Rel | 0.984 |
| dAMP-w | 0.43 |
| Lys-Rel | 0.508 |
| Lys-w | 0.65 |
| Rel-w | 0.19 |

**ns**
