## Extended Material and Methods for "*Drosophila* antimicrobial peptides and lysozymes regulate gut microbiota composition and abundance"

### 16S rRNA gene amplicon sequencing and data processing

Divisive Amplicon Denoising Algorithm 2 (DADA2) pipeline ("dada2" package version 1.14.1 in R) was used to process the sequencing data. All functions were run using the recommended parameters (https://benjjneb.github.io/dada2/tutorial.html) except for "expected errors" during the filtering step which was set to (maxEE=2,5) in "filterAndTrim" function. The RDP database was used for taxonomy assignments. Downstream analyses were performed in R version 3.6.0. Reads belonging to mitochondria, chloroplasts, and eukaryotes were excluded from further analyses ("phyloseq" package version 1.30.0, "subset_taxa" function (80)). Only the variants present in at least 10 samples with a total of 100 reads were retained for downstream analyses ("genefilter" package version 1.68.0, "filterfun_sample" function (81)). To complement the taxonomic classification based on the RDP database, sequence variants were further assigned to the gut microbiota members based on their alignment to the full 16S rRNA gene sequences obtained by Sanger sequencing of each isolate.

### Gnotobiotic fly cultivation and media

To generate germ-free flies, embryos were collected from an overnight egg laying on grape juice-agar plates supplemented with yeast. Embryos were washed with tap water, sterilized by soaking in 3% bleach for 3 minutes and were rinsed with autoclaved water 3 times. ~200 eggs were counted on a mesh under a laminar flow hood and were transferred to filter cap falcon tubes (TPP catalogue #87050) containing the autoclaved larval medium (0.79% Agar, 5.2% cornmeal, 11% sucrose, 4% yeast, 1.12% Moldex, and 0.77% propionic acid, where the last two were added when the medium was <78˚C) supplemented with antibiotics (50µg/µl Ampicilin, 50µg/µl Kanamycin, 10µg/µl Erythromycin, 10µg/µl Tetracyclin).

Upon emergence, 0-2 day old adults were transferred to autoclaved 2% yeast adult medium (all other ingredients were identical to the larval medium) without antibiotics to remain germ-free or to be colonized with commensals. To colonize adults, cultured bacteria were centrifuged at 3000 rpm for 10 minutes and resuspended in sterile PBS to a concentration of ~5 x $10^6$ cells per 100µl. To generate the commensal cocktail, 100µl of each bacterial suspension was mixed in 1:1 ratio to maintain the same cell concentration. 50µl of the bacterial suspension was spread over the adult medium using glass beads (3mm diameter) for 10 seconds and the tubes were air dried under the laminar hood for 2 hours. Germ-free adults raised in antibiotic medium were anaesthetized on ice for 10 minutes, and 20-30 adults were transferred to each tube.
